# SC-Framework: A robust and FAIR semi-interactive environment for single-cell resolution datasets

**DOI:** 10.1101/2025.11.11.687874

**Authors:** Hendrik Schultheis, Jan Detleffsen, René Wiegandt, Mette Bentsen, Yousef Alayoubi, Guilherme Valente, Micha Frederick Keßler, Brenton Bruns, Dlnija Mirza, Angeline Usanayo, Jasmin Walter, Philipp Goymann, Moritz Hobein, Carsten Kuenne, Mario Looso

## Abstract

The accelerated development of single-cell technologies has profoundly impacted the field of biological research, facilitating unparalleled insights into cellular heterogeneity. However, this progress has also produced new computational challenges in the field of bioinformatics: single-cell datasets are increasingly high-dimensional, multimodal, and large-scale, while analysis workflows often remain fragmented, data type specific, ad hoc, and difficult to reproduce. The prevailing methodologies are dependent on a combination of public tools, which hinders the reproducibility of results, limits scalability, and complicates the efforts to establish benchmarks. The necessity for a higher-level, unified framework for single-cell data analysis is paramount to address these inherent limitations. Here, we introduce the SC-Framework, providing the integration of standardized data structures, declarative workflows and standardized computational backends in a containerized environment, enabling analysts to focus on biological interpretation rather than technical overhead. SC-Framework is available at GitHub (https://github.com/loosolab/SC-Framework).

## Introduction

The ever-increasing resolution of sequencing-based methods for various cellular modalities (gene expression, chromatin accessibility, surface protein abundance, etc.), has revolutionized our understanding of molecular processes. Access to the single-cell (SC) level permitted the formulation of entirely new inquiries within the domains of cellular heterogeneity and cell differentiation. However, the increasing volume of data and its integration necessitate sophisticated analysis methods, computational tools, and robust frameworks for data processing. It is important to note that the methods established for bulk sequencing are not uniformly applicable to SC resolution data. While there is an ongoing effort to develop new tools^1^ and an increasing amount of analysis guidelines and best practices workflows^2–5^, a fully automated SC analysis akin to bulk analysis workflows is yet to be achieved.

As of writing, multiple tools aim to provide an environment to simplify and streamline SC analysis (Table 1). Broadly, these tools can be split into four categories: utilities, toolkits, automated workflows and graphical user interface (GUI) workflows.

Each of these has been designed to appeal to a different audience. Utilities aim to solve a specific SC task, e.g. batch correction (harmony^6^) or clustering (leiden^7^). Toolkits provide the individual steps for analysis, making them versatile and flexible, but often provide minimal guidance in the design of a SC analysis (Seurat^8^, Scanpy^9^, EpiScanpy^10^, Muon^11^, squidpy^12^, SnapATAC2^13^, scvi-tools^14^). These toolkits require at least minimal coding experiences that often render them inaccessible for people with a limited computational background. But even experienced developers may run into issues due to lack of standardization considering methods and thresholds in the SC field^15,16^. Toolkits are often aimed at solving higher level tasks and thus either integrate or combine with multiple utilities into a working environment, meaning their computational dependencies have to be managed. This can become challenging, making long-term maintenance cumbersome and significantly hindering portability between systems. Automated workflows (e.g. ENCODE: ATAC^17^, RNA^18^, ChIP^19^, DNA^20^, miRNA^21^ or nf-core pipelines^22^ scnanoseq^23^, scrnaseq^24^) require less individual programming but are static, which means they will always perform the same steps in the same order. While this is beneficial in terms of reproducibility, it becomes an obstacle for data types and analysis steps that rely on dynamic threshold adjustments, as in the case for most SC workflows. GUI workflows (e.g. WASP^25^, GRACE^26^, ASAP^27^, SCALA^28^), often web- and cloud-based services, combine the benefits of flexible workflows with a graphical interface. This enables usage without programming experience and rapid explorative analysis while the impact of thresholds can be shown immediately. The downside of GUI workflows is the limited long-term reproducibility due to lack of automation, as well as missing access to cutting-edge tools since integration is typically time consuming.

Between sequencing and final data interpretation, we have identified two major sections, which we call preprocessing and data analysis. The preprocessing encompasses all steps including the initially sequenced reads, basic quality control (QC) and either cell event identification, or mapping and quantification of the reads. The product (in the case of scRNA) is a matrix defined by read counts in the dimension of cells by genes. While the process of creating the matrix is sufficiently automated^29,30^, working on this matrix is not, and thus the focus of this work. Processing the count matrix mainly includes the interpretation effort of the given data. With SC data complexity, it is crucial to use software that is streamlined for transparency, robustness, and reproducibility while also being extensible to integrate ongoing developments in the field.

Here we present the SC-Framework (Fig. 1). It adheres to the FAIR software principles which is an abbreviation for Findability, Accessibility, Interoperability and Reusability^31^. Following this principle ensures that code and data are well documented, easily discoverable, openly accessible, compatible across systems, and reusable^32^. The overall goal is to maximize understanding and reproducibility of scientific data interpretations. The SC-Framework is designed to minimize technical burden, reducing the workload for researchers performing SC analysis, allowing them to focus on biological interpretation. It is suitable for different audiences ranging from novice users in need of robust guidance to facilities with a large number of projects that require reproducibility, documentation of code, and high throughput.

**Figure 1:**
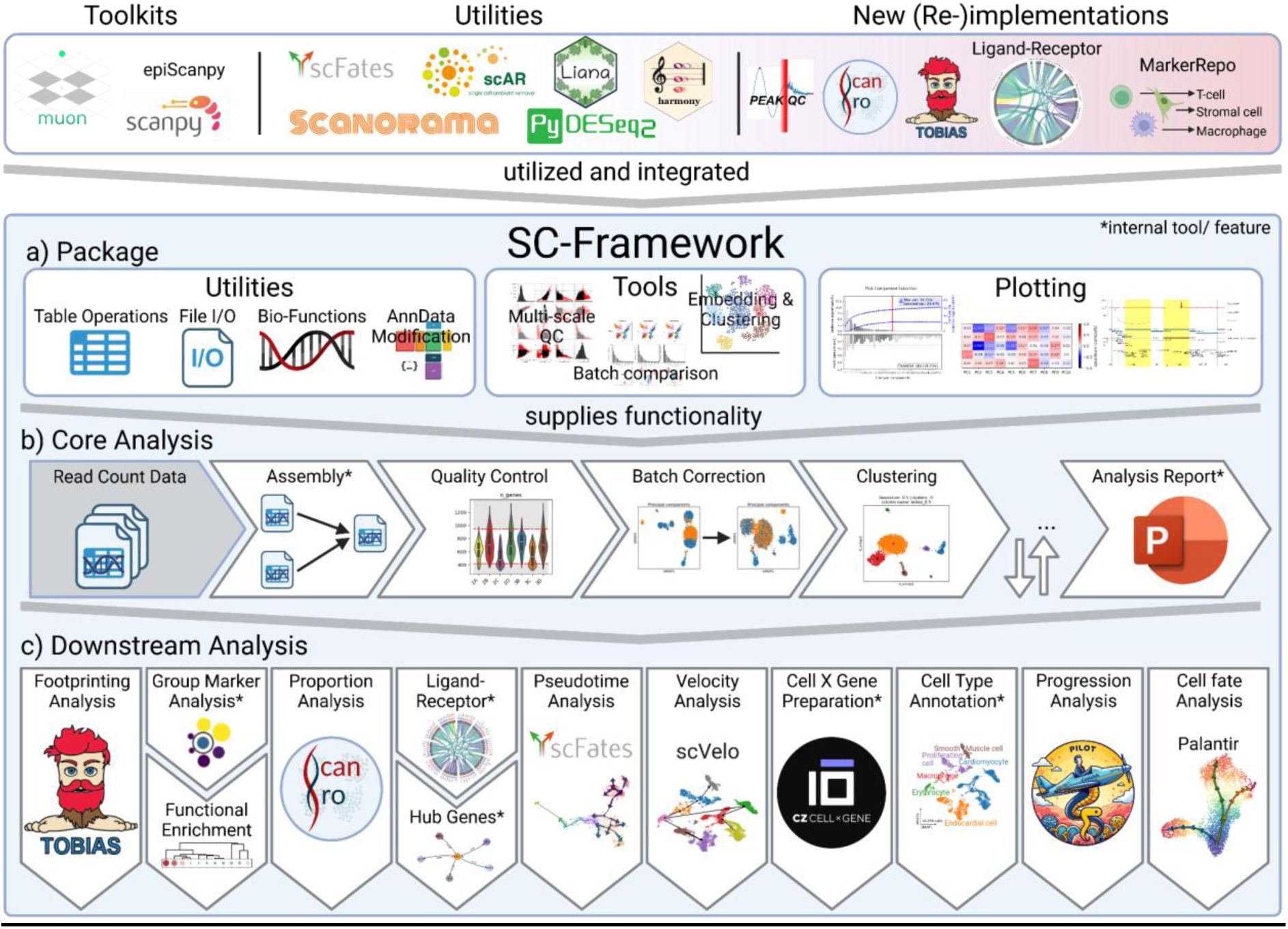
SC-Framework overview. The SC-Framework is an overarching framework for frequently used toolkits, individual purpose SC utilities, and newly implemented SC applications (top). It consists of a Python package (a) providing SC related functionality, and an analysis workflow that utilizes the package to provide a streamlined way to work on SC data (b, c). The workflow is divided into a core analysis part (b) i.e. mandatory steps and downstream analysis (c) i.e. optional steps for individual research focus. Created in BioRender. Schultheis, H. (2025) https://BioRender.com/vm2b5cl

The framework encompasses a collection of utilities and toolkits packaged into an environment (Fig. 1a) and streamlined in a guided step-by-step manner, providing a full analysis workflow for RNA, ATAC, and other modalities. We provide core analysis steps for each modality (QC, batch correction, normalization, and clustering; Fig. 1b) complemented by downstream analysis modules (e.g. annotation, receptor-ligand investigation, proportion analysis) in a highly integrated manner (Fig. 1c). A reporting module collects information such as plots and thresholds to create a comprehensive summary document of the analysis (Fig. 1b). The analysis modules are realized as Jupyter Notebooks, which combine the reproducibility of an automated workflow with the flexibility of a toolkit and the explorative capabilities of a GUI. We capitalize on this versatility with a predefined structure applied to all notebooks. In short, the notebooks are divided into guidelines (explanatory texts, images and tables), highlighted user input fields and immutable code-blocks. This fosters an interpretation-driven environment as it allows users to focus on exploratory rather than computational aspects. It enables corresponding parameter settings to be run interactively, stored and repeated with minimal effort. The automatic generation of logs and a folder structure for intermediate steps foster reproducibility of data interpretations. The SC-Framework produces publication-ready plots and corresponding information to be included in the method section of manuscripts.

## Results

### A FAIR single-cell framework providing a guided analysis

We aligned the SC-Framework with the FAIR principle by organizing it into two main parts, namely i) a Python package that offers encapsulated functionality for working with data at SC resolution, and ii) a series of Jupyter notebooks in a predefined order that leverage the package’s capabilities to provide a flexible scaffold for analysis.

In case of i), the package is an implementation of utilities, tools, and plotting functions from the scVerse such as Scanpy^9^, EpiScanpy^10^, and Muon^11^. It extends existing methods and implements new ones not yet standardized in the SC field by making use of standardized general interfaces within the framework. The package wraps the universal Python AnnData^33^ format for data handling, which allows for storing layered high-dimensional data and corresponding metadata in one type of object. The design of the package is highly modular to allow long-term maintenance, adaptation to new data formats, and universal use of routines to avoid code duplication for related problems within the SC field (e.g. annotation, embedding, filtering).

In case of ii), the series of Jupyter notebooks feature a predefined but flexible and adaptable workflow that constitutes an SC analysis workflow. Each Jupyter notebook represents one major step of the SC analysis, defined by an input data object, a series of analysis steps, and an export object that is used as input for the successor notebook. Overall, our notebooks follow best practices where possible^2,4,5,34,35^, and provide alternative branches of commonly used methods when there is no clear consensus.

For ease of use, each notebook follows a similar structure with recurring structural elements (Supp. Fig. 1). At the top, a highlighted cell indicates the need for user interaction, and is generally intended for setting analysis parameters such as the organism to be studied, thresholds for analysis and various parameters that require user input. By default, most values are preset with defaults. Other cells are not intended for editing by default, to protect novice users from introducing unintended changes. Marker lists for e.g. cell cycle stage prediction, sex chromosome genes, rRNAs and other groups mainly used for various QC steps are dynamically loaded from within the framework. Each notebook features manual blocks providing information on a specific step, links to additional resources, as well as guidance on the interpretation of the data (e.g. the meaning of QC metrics and how a suitable threshold may look like). The SC-Framework includes an introductory video series (https://youtube.com/playlist?list=PLTA07KFyG53ImxJKpoiPO9XiU_I5anMKp&feature=shared) specifically catered to first-time users as an entry to the SC-Framework ecosystem, explaining idioms and structures, as well as guiding users through their first analysis. An extensive manual (https://loosolab.pages.gwdg.de/software/sc_framework/) provides information and examples for advanced users wishing to customize or create new analysis. Unlike most existing workflows (Table 1), the SC-Framework implements self-documenting objects. This feature automatically creates and stores a protocol of the applied analysis steps within the data object. The protocol encompasses details on the called function (e.g. name, timestamp) and its corresponding parameters (filter thresholds, input files, etc.) to follow the analysis process at any given time directly from the data object. This feature seamlessly integrates into the AnnData format^33^. This capability, combined with configuration files that specify notebook order and directory structure, enables iterative analysis, the option to revisit and adjust parameters without losing track of prior steps. And supports branching workflows, such as comparing outcomes of different parameter choices (e.g., batch correction methods) or bypassing computationally intensive steps by diverging from a specific analysis point. (Fig. 2a-c).

**Figure 2:**
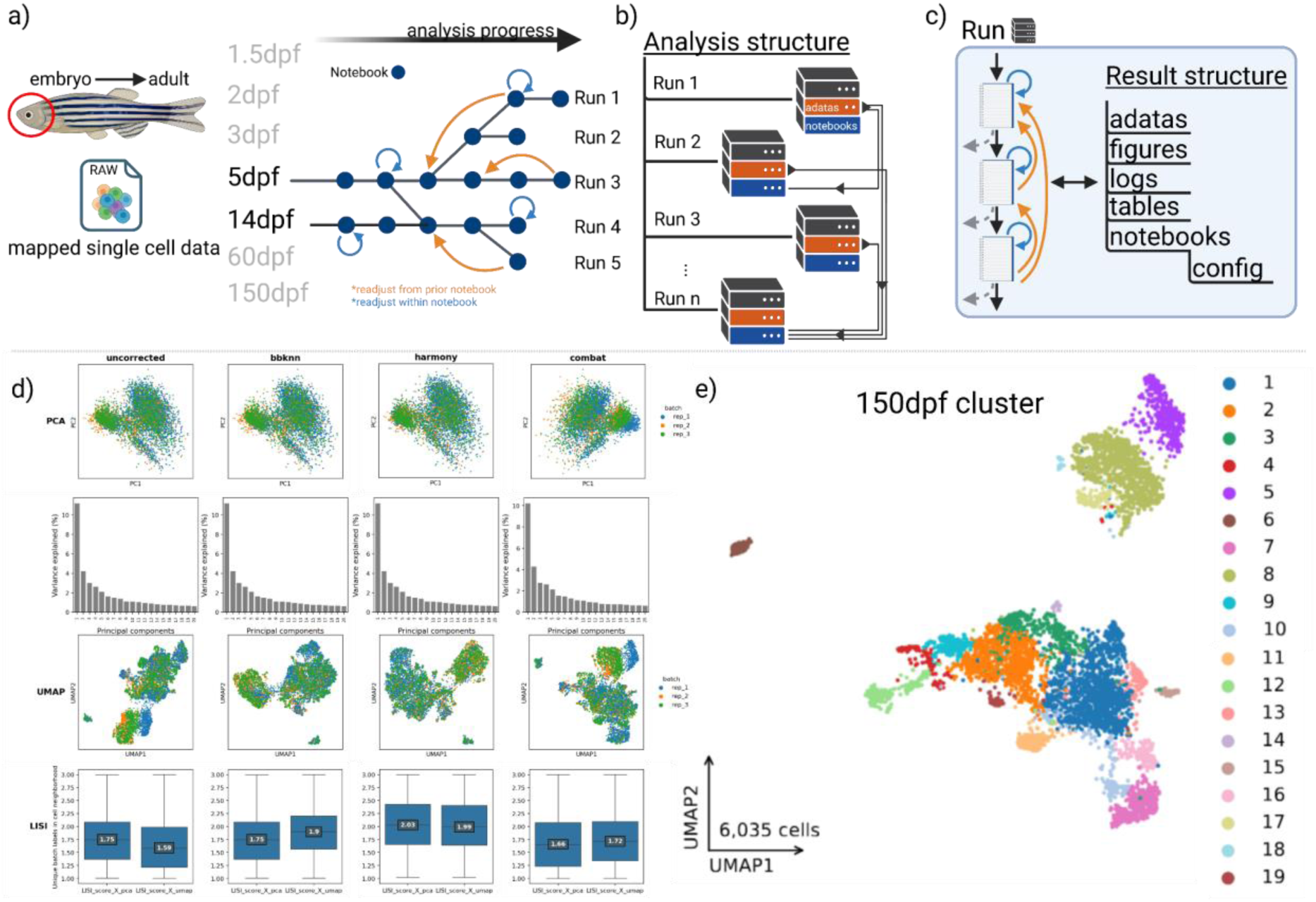
exemplary scRNA core analysis on zebrafish. a) Zebrafish samples from multiple timepoints are processed by the notebooks. The graph illustrates progress of an exemplary analysis, showing branching (two lines exiting or entering a notebook), or looping as blue and orange arrows (see c). b) analysis runs are saved as a stack of directories. AnnData objects may be used in a notebook of a subsequent run indicated by the arrows. c) Scheme of an analysis run. Notebooks are run in order (top to bottom) with the option to rerun a notebook (blue arrow; loop) or restart from a prior notebook (orange arrow; loop), the dashed arrows represent potential branches to another run. Running the notebooks automatically populates the folder structure. d) A grid of embeddings (PCA and UMAP) and their components colored for different metrics over different batch correction tools (columns) and a LISI score plot (bottom row) showing the effect of each tool. e) UMAP of the 150 dpf sample colored by clusters. Created in BioRender. Schultheis, H. (2025) https://BioRender.com/8jj39hf

To simplify reproducibility and interoperability on a higher layer, especially in the context of resource-intensive analysis steps, we consistently apply the concept of application virtualization by providing Docker^36^ containers for each release of our framework that can be easily deployed locally or on a centralized compute cluster. These environments are reproducible units of computational tools and workflows with pinned software versions that can be loaded on demand. In summary, the qualities mentioned above render the SC-Framework a FAIR software, intended to generate reproducible workflows and a documented and FAIR output. At the same time, the package with its notebooks supports intermediate workflow entrance, loops and branching of the workflow to accommodate the flexibility needed for the vibrant SC field.

### Flexible building blocks enable looping and branching within a scRNAseq analysis

The SC-Framework allows for flexible analysis runs, including looping and branching, while ensuring FAIR data handling. Looping and branching are integral parts of most analyses. Nevertheless, their ubiquity can render them inconspicuous, as they naturally occur throughout the analysis. We define looping as re-running parts of the analysis while adjusting parameters. In traditional workflows, this requires repetition of the whole analysis, a tedious and inefficient process. The SC-Framework supports looping back to a previous step at any time, either within a notebook by running one or more previous code blocks, by restarting from the beginning of the current notebook, or by jumping back to a previous notebook and re-executing the analysis (Fig. 2a). Looping is a quick way to try different configurations, therefore, it won’t track anything but the last configuration. Permanent tracking of multiple configurations is meant to be done via branching. We define branching akin to *git branches*, where an analysis can be split into multiple parallel analysis runs, for example, to explore data subsets after QC, or to combine multiple analysis runs, allowing to analyze multiple datasets together. Branches are realized as independent directories. The first notebook in a run folder, the assembly notebook, links to the data of the preceding run (Fig. 2b). Like all notebooks, it accesses a hierarchically organized configuration file (Fig. 2c), harboring global parameters, default values, relative paths and global resources to minimize the overhead to the essentials, while at the same time leaving the flexibility in the hands of experienced users. Branching can be done after each notebook (Fig. 2c). In this section, we showcase this ability and provide an overview of the included features by analyzing an scRNA Zebrafish dataset of the cranial neural crest spanning seven time points encompassing embryonic development to adulthood^37^ (Fig 2a). Of note, we performed the exemplary analysis on the zebrafish data independently for each single timepoint and subsequently merged the individual analysis runs in order to demonstrate the flexibility of the framework and the branching and looping features.

Starting from the raw count matrix containing gene transcript counts per cell, we performed the core analysis which is encapsulated in four modules (Jupyter notebooks, Fig. 1b, Supp. Fig. 1). These four core modules are **i) the assembly, ii) QC, iii) batch correction, and iv) embedding/clustering**.

### Core analysis

The **i) assembly notebook** is intended to reformat the data into the AnnData format^33^ to allow further processing. It is set up to import and handle common SC related formats from the R ecosystem (Seurat^8^, SummarizedExperiment^38^), text-based files, typically consisting of three tables (expression-matrix, barcodes and features, such as often provided by GEO^39^ and other public data repositories), and the Python ecosystem-related AnnData format. Multiple files originating from e.g. samples, conditions, or multiple datasets, are merged into one large object, allowing for flexible integration of arbitrary data points. To account for inconsistencies created during the merging process, such as outdated QC metrics, we added the option to omit or rename variables, observations, and secondary matrix layers. For the zebrafish data this meant combining replicates where applicable (Table 2).

The **ii) QC notebook** removes low-quality cells and genes (accessibility related features in case of ATACseq) based on common metrics (e.g. total counts, mitochondrial content) (Supp. Fig. 2a) proposing either a global or sample specific threshold using the median absolute deviation or a related method using a gaussian mixture model (See methods). Thresholds are customizable by interactive plots. Other commonly used methods, including doublet removal^40–42^, ambient RNA denoising^43^, and the identification of mitochondrial, ribosomal, or gender genes are optionally available.

The **iii) batch correction notebook** is used to adjust the dataset to focus on factors under investigation by providing methods for normalization and exclusion of unwanted variances (e.g. technical variances). Typically, normalization corrects for sequencing depth and outliers. Subsequently, the optimal set of highly variable genes (HVG) is detected by comparing their mean expression and variance-to-mean ratio to reach the optimal range of 1,000 to 5,000 HVGs^5^. To further reduce the dimension of the SC expression matrix, we embed it into a low-dimensional space by computing principal components (PCs) on the HVGs^44^. To find the PCs that reflect primarily cellular expression profiles, we correlate all PCs with given QC metrics calculated earlier, such as total counts, mitochondrial content, or cell cycle phase, and provide a visually supported tool to subset PCs to the biologically most relevant PCs, as well as PCs explaining sufficient variability (Supp. Fig. 2b-c). The final PC selection is then employed to construct a neighbor graph.

The third notebook ends with a comparative batch correction module. Other studies benchmark a selection of datasets to provide a general guideline on the individual tool performance^45^. However, comparison of multiple batch correction methods in the context of a specific dataset would often be preferable. As such, workflows^46,47^ were developed to ensure selection of the optimal batch correction tool for an individual dataset by assessing correction performance. In order to provide this functionality as a fully integrated module, our implementation includes five widely used batch correction tools (BBKNN^48^, MNN^49^, Harmony^6^, Scanorama^50^, and ComBat^51^), as well as integrated plots that illustrate the uncorrected dataset alongside the batch corrected versions (Fig. 2d). In addition, we included the Local Inverse Simpson’s Index^6^ (LISI) which scores the “mixedness” of the batches in the embeddings. In the exemplary dataset, we applied ComBat^51^ to the *150 dpf* zebrafish data shown in Figure 2d-e to integrate three replicates (Table 2).

The final **iv) clustering notebook** features detailed options on computing and fine-tuning embedding and clustering (Fig. 2e). It allows for computing UMAP or t-SNE embeddings. To optimize the overall layout, the chosen embedding is computed in a range of parameter configurations from which the optimal representation can be chosen (Supp. Fig. 2d). Finally, similar to the embedding module, multiple clusterings are calculated and illustrated within the chosen embedding (Supp. Fig. 2e). Modules enable optional refinement, including cluster splitting or merging. (Supp. Fig. 2f). The final clusters are visualized for selected data layers to evaluate the clustering in terms of biological variance and to delimit against technically induced pattern formation (Supp. Fig 2g-h).

In summary, we applied our core notebook stack to the zebrafish cranial neural crest dataset (accessible in detail in the Supplementary Repository) and generated an independent analysis within mere hours. It resulted in a data interpretation closely resembling the publication by Fabian et al.^37^. In line with Fabian et al., we retained 6,035 cells after QC (6,099), which split into 19 clusters (22; Table 2).

#### Downstream analysis

Based on cell identities and their distribution between different conditions, a plethora of tools are available to perform downstream analyses (Table 3). Several integrated downstream analysis notebooks (Fig. 1c) are provided to answer typical biological questions that arise in context of a SC dataset, which in contrast to the core notebooks are not intended to run in a specific order. Most downstream notebooks are applicable to either RNA- or ATAC-derived data, with exceptions like the velocity analysis, which is exclusive for RNA. The downstream notebooks include functionality for proportion analysis, cell type prediction, receptor ligand analysis, marker detection and differential analysis, gene set enrichment methods, trajectory and time line analysis, and export of a h5ad object for optimal representation in the widely used CELLxGENE app^52^, as well as an automated summary report notebook.

To further benchmark our notebook stack, we attempted to reproduce the more detailed analysis from Fabian et al^37^.

The core analysis generates a UMAP embedding and initial clustering, assigning cells to putative functional groups (Fig. 2e). We evaluate whether the “group_marker” notebook, which was designed to identify cluster-specific features such as marker genes, can recover well-established cell-type-specific markers. Assessing markers across multiple clustering resolutions aids in refining and validating cell groupings.

The initial clustering showed a suboptimal marker panel with 21 clusters (Supp. Fig 2e, Supp. Fig. 3a, c). Looping back to the clustering notebook, we split and recombined to 19 clusters (Fig. 2e, Supp. Fig. 3b), which improved the visual separation of clusters within the UMAP (see the supplementary repository for details), as well as in the subsequently repeated marker detection (Supp. Fig. 3c, d). For example, the gene *ucmaa* was described to be primarily expressed in cartilage^53^, the fibroblast markers *hpdb, hgd*^37,54^, and the muscle-related *thbs4b*^55^ and *myoc*^56^, are all found within the top 15 markers of their respective clusters.

**Figure 3:**
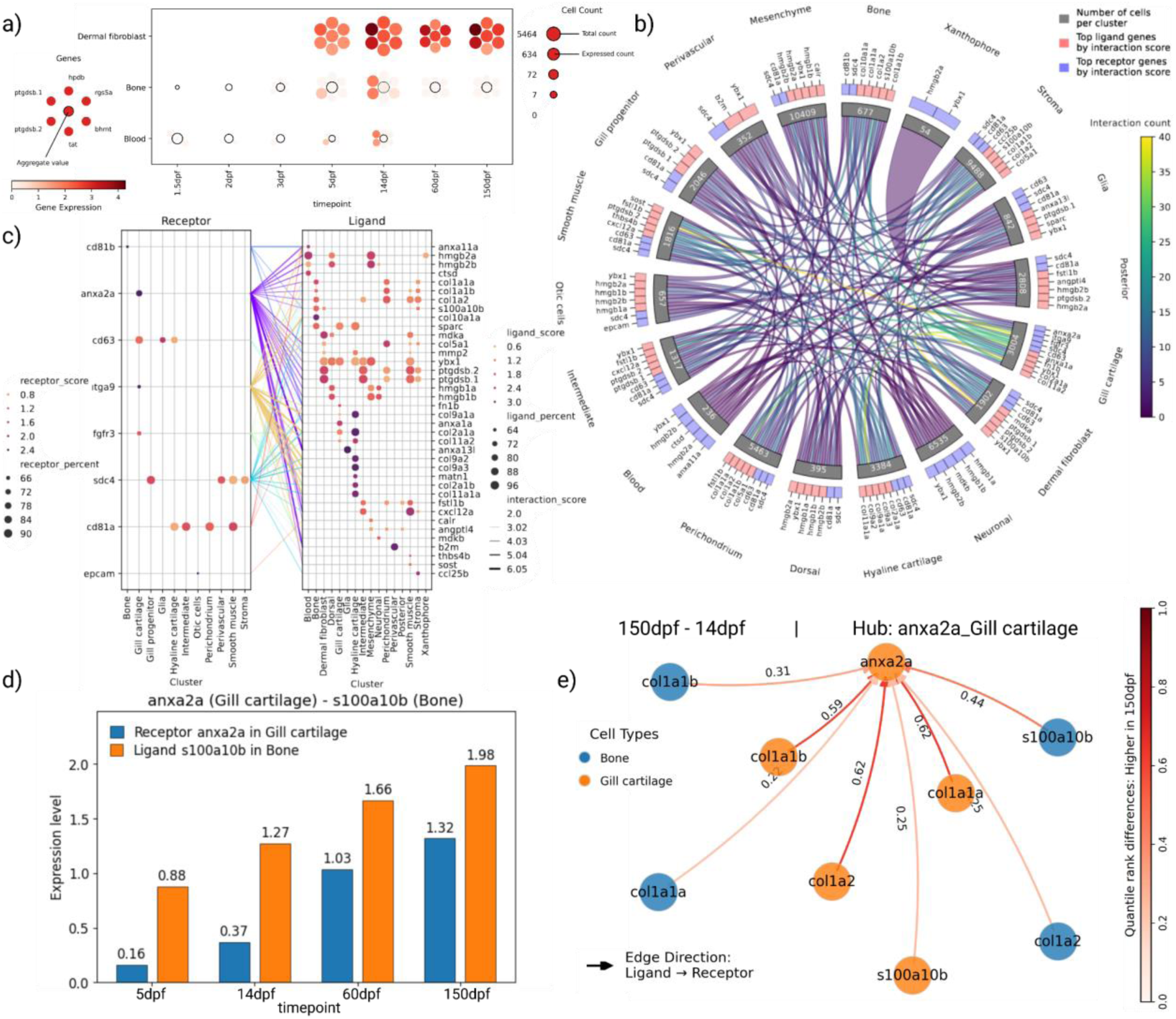
exemplary scRNA downstream analysis of the combined timepoints on zebrafish. a) A **planet plot** showing the gene markers (planets/dots) for a subset of cell types (rows) across timepoints (columns). The central dot per planet system indicates the mean expression of the moons. b) **Cordplot** quantitative interactions between zebrafish cell types. The outermost shell shows top receptor and ligand genes per cell type. c) **Focused interaction plot** presenting a qualitative view on most prominent receptors (left) connected to their respective ligand (right) between cell types. d) **Single interaction plot** for one receptor-ligand pair between specific cell types over time. e) **Hub plot** illustrating quantile rank differences of receptor-ligand interactions between 14dpf and 150dpf, with genes and clusters selected from c). Each node represents a gene (receptor or ligand), with interactions and their difference between timepoints shown as a colored arrow. Created in BioRender. Schultheis, H. (2025) https://BioRender.com/f1rl8cl

The SC-Framework supports marker identification via either the Scanpy^9^ provided functionality based on statistical tests (e.g. Wilcoxon) of a dedicated cluster vs. all remaining cells, or a module encapsulating DESeq2^57^ for pairwise cluster or condition comparisons. The notebook is complemented by various plotting options, most notably the marker planet plot (Fig 3a). This plot is intended to visualize differences in conditions and groups by incorporating selected features and a summarized score. Identified markers are valuable for gene set enrichment analysis (GSEA) or for annotation.

Cell type annotation represents a critical step of SC analysis and typically involves computational approaches, such as reference-based (e.g. SingleR^58^ or scmap^59^), marker-based (e.g. CellAssign^60^ or SCSA^61^), or machine learning based methods (e.g. SingleCellNet^62^ or TOSICA^63^). These are typically complemented by expert knowledge and manual curation. SC Framework supports annotation via SCSA and MarkerRepo. The MarkerRepo implements a reference-based approach, with an interface that facilitates importing, storing, and managing marker lists with optional metadata (e.g. tissue, age, organism) from databases or publications. Two scoring methods are available to weight marker genes/peaks during annotation. The first is based on the frequency over all selected groups (e.g. cell types), and the second is the ubiquitousness index (UI), such as implemented in the PanglaoDB^64^. Of note, PanglaoDB and other reference databases are restricted to mostly human and mouse genes. Therefore, the MarkerRepo implements additional functionality to automatically transfer marker lists to other species of interest based on a homology database (BioMart^65^ or HomoloGene^66^), enabling the annotation of any organism. The annotation notebook is geared to streamline the process of searching and combining marker lists, either supplied by the user or already included within its database. In the case of the 150 dpf zebrafish cranial neural crest dataset, we collected the corresponding markers reported by Fabian et al. (Non-mesenchymal CNCC derived, Non-CNCC, 150 dpf), and submitted them to the MarkerRepo for annotation. The annotation notebook optionally runs all algorithms and scoring schemes in parallel including comparison to an optional reference. In the exemplary 150dpf dataset, we used the annotation provided by Fabian et al.^37^ as a reference (Supp. Fig. 3b, Table 2).

Finally, the annotated dataset is stored within the SC-Framework folder structure (Fig. 2b-c) to become accessible to other notebooks. Here, we looped back to the marker notebook, since the cell type annotation combined clusters by assigning them to the same cell type (Supp. Fig 3b). We then applied the GSEA notebook to find and visualize gene ontology (GO) terms enriched within marker sets. As expected, we found GO-terms matching the annotated cell types with genes of the *phenylalanine catabolic process* enriched for the *dermal fibroblasts*, as reported by Fabian et al. (Supp. Fig. 3e, f).

We performed individual analyses of all other timepoints similar to 150dpf (Fig. 2a). In order to demonstrate branching and joining of workflows (Fig. 2a-c), we combined the resulting data objects with the assembly notebook to identify developmental trends during the maturation of zebrafish. Next, we calculated an integrated embedding and clustering via the normalization and clustering notebook for the combined dataset, leaving out QC and batch correction attempts. The latter was not applicable, since biological variance correlated with technical batches for this dataset, but is typically necessary.

Clusters were refined by observing markers, followed by annotation with the combined marker lists of Fabian et al.^37^. The annotated embedding of combined timepoints was used to investigate developmental changes, particularly the emergence of specific cell types. To answer this question, we utilized the proportion analysis^67^ notebook to calculate significant changes for each cell type per timepoint (Supp. Fig. 3g). In agreement with Fabian et al. we found cell types emerging at specific timepoints, such as *dermal fibroblasts* and *bone* in larvae and later stages.

Finally, we applied our receptor-ligand (RL) analysis to the dataset (see methods), which is intended to predict cell communication between cell types. The notebook provides visualization emphasizing global aspects (e.g. which cell types are highly interactive, supported by a high number of expressed RL pairs (Fig 3b), and individual RL interactions showing high activity between >2 cell types (Fig 3c).

For instance, we found the cell type *gill cartilage* to be highly interactive with most cell types in the dataset (Fig. 3b). Upon closer inspection, we found *anxa2a* to be a key gene for these interactions, well known to be involved in processes such as regeneration^68–70^, development^71,72^ and immune responses^73^, targeted by various ligands from different cell types. We defined such genes as hub genes and examined potential changes in the cellular communication along the timeline. By utilizing the hub notebook, we investigated differences in interactions, and found the *anxa2a* hub in *gill cartilage* to follow the expression pattern of its ligand *s100a10b* in *bone* (Fig. 3d, e). Human orthologs of *anxa2a* and *s100a10b* (*AnxA2, S100A10*) were described to form a complex involved in important biological processes such as exocytosis and membrane repair^74^. Both genes were suggested to have a function in matrix vesicle-mediated mineralization, a process needed for bone formation^75^.

In summary, the iterative steps described above underscore the significance of a versatile and resilient workflow, facilitating looping or branching within a robust framework, thereby ensuring the availability of data layers that are essential prerequisites for subsequent analysis steps by deep integration of publicly available and internally implemented tools.

### Generic implementation enables the analysis of different modalities

The SC-Framework is designed and implemented by a generalized approach to allow the reuse of large code blocks, modules and wrappers. This design supports the application to a wide range of datasets and data types, as well as the fast adaptation and generation of notebooks to new data types. The notebooks intensively reuse the generic code, and the group of downstream notebooks are prone to be used with multiple data types, with no need for modification. However, some data types such as scATACseq need data-specific operations.

In order to demonstrate specific steps in this workflow and the types of analysis that are integrated into the framework, we use an exemplary single-nucleusATAC (snATAC) data of human peripheral blood mononuclear cells (PBMCs) from a healthy donor, obtained from 10x Genomics^76^. At a higher level, the steps of the ATAC workflow follow a similar structure compared to its analog on RNA analysis. The first four notebooks are the core steps, including a) the assembly, i.e. converting and preparing the dataset for the following analysis; b) QC steps, which lack widely accepted concepts; c) batch correction; and d) embedding and clustering.

In case of b) it is noteworthy that we offer a series of metrics to filter cells. In detail, we integrated the ability to use ATAC-seq specific modalities as quality proxy, like the fragment length distribution (FLD)^77^, the amount of fragments overlapping with regions of interest e.g. promoters, the fraction of reads in peaks (FRiP), and the transcription start site enrichment (TSSE). Scores that rely on external information e.g. fragment locations are optional and will be skipped or generated depending on the respective information availability. We provide integrated peak annotation to nearby genes by distance^78^, which enables gene-related downstream analysis such as gene set enrichment or cell type annotation. A notable step of the QC notebook is the binarization of the feature matrix. The chromatin state is often described in a binary notion, as either “open” or “closed”. This has the benefit of negating potential sequencing depth-related issues^79^. However, this choice is under debate, as recent studies show that binarization can result in a loss of information as it disregards chromatin differences between alleles ^4,79,80^. We acknowledge this ongoing debate by making binarization optional.

One of the next steps is normalization, where we provide the widely used method Term Frequency-Inverse Document Frequencies (TF-IDF), which aims to highlight important features based on their occurrence in the dataset. However, recent studies showed that this method should not be treated as a depth-normalization technique but rather as a weighting scheme. As such, it is suitable for clustering but may not apply to other analysis steps. The increased sparsity of scATAC influences differences in sequencing depth, which is exacerbated by TF-IDF^79^. As a direct result, one dimension reduction component correlates with the total counts. The recommended approach for this issue is to remove the offending component (typically the first one)^81^. In our workflow, we visualize and provide a filtering module for dimension reduction components by calculating the correlation of a variety of QC metrics (including total counts, Supp. Fig. 2c). Of note, it recommends a subset of components for subsequent analysis steps.

In case of c), we skipped batch correction as the experimental design did not allow us to discriminate between biological variance as the dataset contains only one sample/ condition (healthy). Consequently, we continued with d) to generate an optimal embedding and clustering (Supp. Fig. 4a). This concludes the core analysis.

Next, we aimed to identify cell types to investigate differential changes in transcription factor binding. Genes of the previously annotated peaks are investigated to identify group markers utilizing the marker notebook as described above for the RNA dataset. The resulting marker genes for each cell cluster were then used to annotate cell types using the annotation notebook (Supp. Fig. 4b). As expected, and described by Kleiveland^82^, we found the PBMC cell types. The amount of each of the cell types matched the expected composition described in the MACS handbook^83^, except for the Dendritic cells, which show a higher-than-expected content (Supp. Fig. 4c). The increase might be explained by the annotation combining *Dendritic cells* and *Monocytes* due to their shared lineage.

Next, a differential transcription factor binding analysis is realized using scTOBIAS^84^ available through a notebook of the same name. Cell signals from user defined groups (e.g. cell type + specific condition) are exported to pseudo bulk cell groups and send to an integrated TOBIAS wrapper that performs selected pairwise comparisons on transcription factor binding occupancy. Analysis of the PBMC cell types between *Naive CD8^+^ T cells* and *Regulatory T (Treg) cell*s, a derivative of *CD4^+^ T cells*, identified transcription factors, in line with expectations. For example, BACH2^85^, BATF^86^, JUNB^85,86^, BATF3^87^, SATB1^88,89^, NFIL3^90^, ATF4^91^, BACH1^92^ are associated with *CD4^+^ T cells* and EGR1^93^, EGR2^94^, KLF2^95^, KLF4^96^, SP1^97^, Nrf1^97^, Ahr::Arnt^98^ are associated with *CD8^+^ T cells* (Supp. Fig. 4d). While most downstream notebooks are agnostic to the data, meaning they can be used for both chromatin and transcriptome analysis, the TOBIAS^84^ notebook is unique to ATAC.

In summary, our ATAC notebooks indicate the generalized design and concept of our code, seamlessly integrating various data types and projects of different size/ cell numbers. The notebook further ensures seamless integration into the folder structure of the framework supporting findability and reproducibility by storing in- and outputs in a comprehensive manner. We have used our notebooks to provide access to datasets including scRNAseq, multimodal data, scATAC, CITEseq, spatial scRNAseq, Immune Panel Seq and others, from which a series was already published in peer reviewed journals (Tombor et al.^99^, Khassafi et al.^100^, Dizdarevic et al.^101^, Panza et al.^102^, Xu et al.^103^, Wang et al.^104^, Goumenaki et al.^105^, Malainou et al.^106^, Jang et al.^107^, Da Silva et al.^108^, Cardeira-da-Silva et al.^109^, Juan et al.^110^, Roquid et al.^111^, Chu et al.^112^, Detleffsen et al.^77^).

## Discussion

SC analysis is a complex topic often perceived as a convoluted process of opaque instructions, worsened by the fact that a substantial part of the analysis steps do not have a gold standard solution, thus requiring ad hoc decisions based on the dataset, which rely on the analyst’s experience. Finding solutions is tied to exploring the data, which needs programming knowledge to use the full repertoire of state of the art tools, thereby decreasing the accessibility of SC analysis. Conducting SC analysis is thus a highly skilled task that requires an analyst with extensive knowledge in programming, biology and sequencing especially focused on SC applications. These issues hinder reproducibility of data interpretations, as small deviations in the order of steps and thresholds can have a profound impact on the outcome.

While progress has been made to open analysis to a wider audience, very few toolkits follow a concept that aims to fulfill the FAIR principles and simultaneously keeps the flexibility that is currently needed for an explorative analysis. For example, best practice resources are a great introduction to the SC topic, however they often lack details when it comes to specific considerations during analysis and are not directly applicable to a dataset. On the other end, automated workflows are ready-to-use and reproducible, but they lack the flexibility needed for data exploration, especially within the workflow. The SC-Framework aims to combine the advantages of both approaches by utilizing a python package in combination with Jupyter Notebooks. This allows us to i) define the analysis steps with minimal coding requirement ii) add descriptions and references to each step, iii) retaining flexibility through allowing on-demand creation of new steps, while iv) storing the Jupyter notebooks, with respective data files in a comprehensive folder structure, and v) automatic logging to ensure reproducibility and findability.

To expand the audience of the framework from computational analysts and professional facilities to non-computational biologists, we spent considerable efforts to provide step by step manuals and a series of videos spanning the whole project from introduction to downstream analysis, including a full API documentation and coding examples for most functions. Our Continuous Integration (CI) pipeline ensures robustness of the code base through extensive unit tests and provides Docker based containers for each version to guarantee reproducibility of past analysis.

Our integrated report generator allows the analyst to produce and share a well-structured PowerPoint of the current analysis state, supporting the often-overlooked communication between analysts and persons interested in the data interpretation, if not the same individual. In summary, the SC-Framework provides a centrally maintained, comprehensive and fully guided SC analysis environment catering to novice and experienced users alike. Provided modules cover a wide range of types of analysis typically applied to SC resolution datasets. Its flexible and generalized approach allows for the extensibility needed to stay relevant in a fast-paced field such as SC.

## Data Availability

Quantified zebrafish cranial scRNA data of Fabian et al.^37^ was downloaded from GEO accession GSE178969. Human 10k healthy PBMCs used for snATAC analysis are downloaded from 10x Genomics (https://www.10xgenomics.com/datasets/10k-human-pbmcs-atac-v2-chromium-controller-2-standard). See our GitHub repository for detailed information on the conducted analysis (https://github.com/loosolab/Code_Schultheis_et_al_2025_SC-Framework).

## Methods

### SC-Framework

The SC-Framework is an environment to facilitate full SC analysis starting from quantification data (raw counts) over embedding and clustering to downstream analysis such as cell type annotation or marker identification. It is split into a Python package, providing functionality, and a collection of Jupyter notebooks, detailing the analysis. The SC-Framework with all its resources (documentation, API, etc.) can be accessed in our GitHub (https://github.com/loosolab/SC-Framework).

### The Python package - Sctoolbox

The sctoolbox is a Python package providing the bulk of the SC-Frameworks functionality. It can be installed either through GitHub or the Python Package Index (PyPI). We ensure code quality standards by continuously applying style and type checks as well as unit testing. Likewise, we apply semantic versioning and maintain a changelog to enhance reproducibility and long-term usage. The package provides a full API documentation including example snippets and respective outputs, and references where applicable.

The sctoolbox features two different logging systems. The traditional logging of custom status messages in different severity levels (info, warning, error, etc.), can be written to a file, directly displayed in the console, or both. The second system is function-level logging which automatically stores function calls and their parameters in the AnnData.uns attribute of the AnnData object, making the object self-documenting and all analysis steps traceable. This is realized using Python Decorators, allowing for an effortless inclusion of new functions (*sctoolbox.utils.decorator.log_anndata*).

The package provides automatic threshold (*sctoolbox.tools.qc_filter.automatic_thresholds*) calculations on cells (AnnData.obs) or features (AnnData.var). Threshold calculation can either be done using the commonly used median absolute deviation (MAD) with independent upper and lower bounds 𝑀𝐴𝐷_𝑠𝑐𝑜𝑟𝑒_ = 𝑚𝑒𝑑𝑖𝑎𝑛(𝑣) ± 𝑀𝐴𝐷 × 𝑛_1,2_, or a similar function that uses the standard deviation of the largest Gaussian mixture model component 𝑐 = 𝑙𝑎𝑟𝑔𝑒𝑠𝑡𝑐𝑜𝑚𝑝𝑜𝑛𝑒𝑛𝑡(𝑣), 𝐺𝑀𝑀_𝑠𝑐𝑜𝑟𝑒_ = 𝑚𝑒𝑎𝑛(𝑐) ± 𝜎_𝑐_ × 𝑛_1,2_. The function also accepts external threshold functions.

### The analysis notebooks

The SC-Framework features a collection of analysis Jupyter notebooks, which can be found in our GitHub repository (https://github.com/loosolab/SC-Framework). Method specific notebooks, e.g. RNA related, are found within the “*_analysis” folder, while notebooks usable with any data are found in the “general_notebooks” folder. An analysis starts with copying the “*_analysis” to the desired location and opening the first notebook (assembly). Depending on the circumstances (e.g. whether the data is pre-analyzed) the other notebooks are either run in the order indicated by the number prefix in their name or skipped. Notebooks without a prefix (general) or with a prefix that contains a character (e.g. “0A1_…”) are considered downstream and to be run after the numbered notebooks. While downstream notebooks don’t follow an overarching order of execution, notebooks with the same character in their prefix must be run in order, e.g. “0A1_…” -> “0A2_…”. General notebooks should be copied into the respective analysis folder before their execution.

Regarding the order within the notebooks we aim to follow established protocol where possible or provide descriptions for a user based decision. For example, while there are recommendations to compute HVGs after batch correction^5^, other sources stress that the impact of batch correction on expression-level analysis is unclear and that HVGs improve the performance of batch correction^46,47^, which is why we decided to do HVG calculation before batch correction.

### MarkerRepo

The MarkerRepo is a reference annotation tool, implemented within the sctoolbox, to manage genes and other types of markers. It can be accessed using our GitLab (https://gitlab.gwdg.de/loosolab/software/annotate_by_marker_and_features). At its core is the annotation of cells to arbitrary groups, typically cell types, based on gene marker lists. It provides three algorithms for annotation (SCSA, MR score, UI) and includes automatic homology-based gene conversion. The MarkerRepo further contains a database backend, allowing users to easily integrate new marker lists. The database also enables searching and selecting lists based on metainformation, for example, tissue, to create suitable on-demand marker lists collected from the database.

### Receptor Ligand analysis

The RL analysis requires a database with receptor and ligand pairings to proceed. We integrated the LIANA+^113^ package to provide multiple databases of different organisms. A custom database, in the form of a table with a column of receptor-gene names and a column of corresponding ligand-gene names, may be used as an alternative. Interactions are scored for their combined receptor and ligand enrichment between clusters of cells adjusted for the cluster size and scaled for the number of cells expressing the respective gene. The interaction score of a specific interaction is calculated as the sum of the participating receptor and ligand gene scores. The gene scores are calculated as the z-score over the cluster mean expressions, multiplied by the proportion of cells assigned to the respective cluster, and multiplied by the proportion of cells within the cluster expressing the gene. The interaction score is based on the 𝑤_2_-score of Brickman et al.^114^. Interaction scores >0 can be interpreted as “enriched”, while <0 means “depleted”.

The identification of changing interactions between conditions is investigated through the quantile-ranked differences. This is realized by calculating the interaction scores for each condition by combining and ranking the interactions. The difference in rank between two conditions yields the final score for every interaction. Conditions are implemented to be chosen arbitrarily and can be nested. For example, the data may be separated into wild type and multiple treatments and on a second level subdivided into patients. It is also possible to define an order for conditions with an inherent sequence, such as timepoints, to restrict the difference comparisons to the previous and directly following condition.

## Acknowledgments

Funding

This work was supported by the DFG (Deutsche Forschungsgemeinschaft) under Grand (ExStra EXC2026 Translational Hub 2, KFO309 Z Project) to M.L., Hessisches Ministerium für Wissenschaft und Kunst (LOEWE iCANx Z Project) to M.L., DZHK Rhein Main Site, and the Max-Planck Society.

## Author contributions

M.L. conceived the study. M.L., H.S., J.D., R.W. and M.B. designed the software. H.S, J.D., R.W., M.B., Y.A., G.V., M.K., B.B., D.M., A.U., J.W., P.G., M.H. and C.K. implemented the software. H.S. preprocessed and analyzed the data. H.S., J.D., R.W., C.K. and M.L. wrote the manuscript. M.L. supervised the project.

## References

1. Zappia, L. & Theis, F. J. Over 1000 tools reveal trends in the single-cell RNA-seq analysis landscape. Genome Biol. 22, 301 (2021).

2. Amezquita, R. A. et al. Orchestrating single-cell analysis with Bioconductor. Nat. Methods 17, 137–145 (2020).

3. Grones, C. et al. Best practices for the execution, analysis, and data storage of plant single-cell/nucleus transcriptomics. Plant Cell 36, 812–828 (2024).

4. Heumos, L. et al. Best practices for single-cell analysis across modalities. Nat. Rev. Genet. 24, 550–572 (2023).

5. Luecken, M. D. & Theis, F. J. Current best practices in single-cell RNA-seq analysis: a tutorial. Mol. Syst. Biol. 15, e8746 (2019).

6. Korsunsky, I. et al. Fast, sensitive and accurate integration of single-cell data with Harmony. Nat. Methods 16, 1289–1296 (2019).

7. Traag, V. A., Waltman, L. & van Eck, N. J. From Louvain to Leiden: guaranteeing well-connected communities. Sci. Rep. 9, 5233 (2019).

8. Hao, Y. et al. Dictionary learning for integrative, multimodal and scalable single-cell analysis. Nat. Biotechnol. 42, 293–304 (2024).

9. Wolf, F. A., Angerer, P. & Theis, F. J. SCANPY: large-scale single-cell gene expression data analysis. Genome Biol. 19, 15 (2018).

10. Danese, A. et al. EpiScanpy: integrated single-cell epigenomic analysis. Nat. Commun. 12, 5228 (2021).

11. Bredikhin, D., Kats, I. & Stegle, O. MUON: multimodal omics analysis framework. Genome Biol. 23, 42 (2022).

12. Palla, G. et al. Squidpy: a scalable framework for spatial omics analysis. Nat. Methods 19, 171–178 (2022).

13. Zhang, K., Zemke, N. R., Armand, E. J. & Ren, B. A fast, scalable and versatile tool for analysis of single-cell omics data. Nat. Methods 21, 217–227 (2024).

14. Gayoso, A. et al. A Python library for probabilistic analysis of single-cell omics data. Nat. Biotechnol. 40, 163–166 (2022).

15. Adil, A., Kumar, V., Jan, A. T. & Asger, M. Single-Cell Transcriptomics: Current Methods and Challenges in Data Acquisition and Analysis. Front. Neurosci. 15, (2021).

16. Lu, J. et al. scRNA-seq data analysis method to improve analysis performance. IET Nanobiotechnol. 17, 246–256 (2023).

17. Lee, J., et al. kundajelab/atac_dnase_pipelines: 0.3.3. Zenodo 10.5281/zenodo.211733 (2016).

18. ENCODE-DCC/rna-seq-pipeline. ENCODE DCC (2025).

19. ENCODE-DCC/chip-seq-pipeline2. ENCODE DCC (2025).

20. ENCODE-DCC/dnase_pipeline. ENCODE DCC (2024).

21. ENCODE-DCC/mirna-seq-pipeline. ENCODE DCC (2024).

22. Ewels, P. A. et al. The nf-core framework for community-curated bioinformatics pipelines. Nat. Biotechnol. 38, 276–278 (2020).

23. Trull, A., Community, N.-C., Worthey, E. A. & Ianov, L. scnanoseq: an nf-core pipeline for Oxford Nanopore single-cell RNA-sequencing. 2025.04.08.647887 Preprint at 10.1101/2025.04.08.647887 (2025).

24. Almeida, F. M. de et al. nf-core/scrnaseq: 4.0.0. Zenodo 10.5281/zenodo.15004569 (2025).

25. Hoek, A. et al. WASP: a versatile, web-accessible single cell RNA-Seq processing platform. BMC Genomics 22, 195 (2021).

26. Hao Yu, Yuqing Wang, Xi Zhang, & Zheng Wang. GRACE: a comprehensive web-based platform for integrative single-cell transcriptome analysis. 5, (2022).

27. Gardeux, V., David, F. P. A., Shajkofci, A., Schwalie, P. C. & Deplancke, B. ASAP: a web-based platform for the analysis and interactive visualization of single-cell RNA-seq data. Bioinformatics 33, 3123–3125 (2017).

28. Christos Tzaferis et al. SCALA: A complete solution for multimodal analysis of single-cell Next Generation Sequencing data. 21, 5382–5393 (2023).

29. Kaminow, B., Yunusov, D. & Dobin, A. STARsolo: accurate, fast and versatile mapping/quantification of single-cell and single-nucleus RNA-seq data. 2021.05.05.442755 Preprint at 10.1101/2021.05.05.442755 (2021).

30. Zheng, G. X. Y. et al. Massively parallel digital transcriptional profiling of single cells. Nat. Commun. 8, 14049 (2017).

31. Wilkinson, M. D. et al. The FAIR Guiding Principles for scientific data management and stewardship. Sci. Data 3, 160018 (2016).

32. Barker, M. et al. Introducing the FAIR Principles for research software. Sci. Data 9, 622 (2022).

33. Virshup, I., Rybakov, S., Theis, F. J., Angerer, P. & Wolf, F. A. anndata: Access and store annotated data matrices. J. Open Source Softw. 9, 4371 (2024).

34. Jovic, D. et al. Single-cell RNA sequencing technologies and applications: A brief overview. Clin. Transl. Med. 12, e694 (2022).

35. Lafzi, A., Moutinho, C., Picelli, S. & Heyn, H. Tutorial: guidelines for the experimental design of single-cell RNA sequencing studies. Nat. Protoc. 13, 2742–2757 (2018).

36. Merkel, D. Docker: lightweight Linux containers for consistent development and deployment. Linux J 2014, 2:2 (2014).

37. Fabian, P. et al. Lifelong single-cell profiling of cranial neural crest diversification in zebrafish. Nat. Commun. 13, 13 (2022).

38. SummarizedExperiment. Bioconductor http://bioconductor.org/packages/SummarizedExperiment/.

39. Barrett, T. et al. NCBI GEO: archive for functional genomics data sets—update. Nucleic Acids Res. 41, D991–D995 (2013).

40. Xi, N. M. & Li, J. J. Benchmarking Computational Doublet-Detection Methods for Single-Cell RNA Sequencing Data. Cell Syst. 12, 176–194.e6 (2021).

41. Thibodeau, A. et al. AMULET: a novel read count-based method for effective multiplet detection from single nucleus ATAC-seq data. Genome Biol. 22, 252 (2021).

42. Wolock, S. L., Lopez, R. & Klein, A. M. Scrublet: Computational Identification of Cell Doublets in Single-Cell Transcriptomic Data. Cell Syst. 8, 281–291.e9 (2019).

43. Sheng, C. et al. Probabilistic machine learning ensures accurate ambient denoising in droplet-based single-cell omics. 2022.01.14.476312 Preprint at 10.1101/2022.01.14.476312 (2022).

44. Heimberg, G., Bhatnagar, R., El-Samad, H. & Thomson, M. Low Dimensionality in Gene Expression Data Enables the Accurate Extraction of Transcriptional Programs from Shallow Sequencing. Cell Syst. 2, 239–250 (2016).

45. Tran, H. T. N. et al. A benchmark of batch-effect correction methods for single-cell RNA sequencing data. Genome Biol. 21, 12 (2020).

46. Luecken, M. D. et al. Benchmarking atlas-level data integration in single-cell genomics. Nat. Methods 19, 41–50 (2022).

47. Chazarra-Gil, R., van Dongen, S., Kiselev, V. Y. & Hemberg, M. Flexible comparison of batch correction methods for single-cell RNA-seq using BatchBench. Nucleic Acids Res. 49, e42 (2021).

48. Polański, K. et al. BBKNN: fast batch alignment of single cell transcriptomes. Bioinformatics 36, 964–965 (2020).

49. Haghverdi, L., Lun, A. T. L., Morgan, M. D. & Marioni, J. C. Batch effects in single-cell RNA-sequencing data are corrected by matching mutual nearest neighbors. Nat. Biotechnol. 36, 421–427 (2018).

50. Hie, B., Bryson, B. & Berger, B. Efficient integration of heterogeneous single-cell transcriptomes using Scanorama. Nat. Biotechnol. 37, 685–691 (2019).

51. Leek, J. T., Johnson, W. E., Parker, H. S., Jaffe, A. E. & Storey, J. D. The sva package for removing batch effects and other unwanted variation in high-throughput experiments. Bioinformatics 28, 882–883 (2012).

52. Megill, C. et al. cellxgene: a performant, scalable exploration platform for high dimensional sparse matrices. 2021.04.05.438318 Preprint at 10.1101/2021.04.05.438318 (2021).

53. Neacsu, C. D. et al. Ucmaa (Grp-2) is required for zebrafish skeletal development. Evidence for a functional role of its glutamate γ-carboxylation. Matrix Biol. 30, 369–378 (2011).

54. Rajan, A. M. et al. Single-cell analysis reveals distinct fibroblast plasticity during tenocyte regeneration in zebrafish. Sci. Adv. 9, eadi5771 (2023).

55. Subramanian, A. & Schilling, T. F. Thrombospondin-4 controls matrix assembly during development and repair of myotendinous junctions. eLife 3, e02372 (2014).

56. Whitesell, T. R., et al. *foxc1* is required for embryonic head vascular smooth muscle differentiation in zebrafish. Dev. Biol. 453, 34–47 (2019).

57. Love, M. I., Huber, W. & Anders, S. Moderated estimation of fold change and dispersion for RNA-seq data with DESeq2. Genome Biol. 15, 550 (2014).

58. Aran, D. et al. Reference-based analysis of lung single-cell sequencing reveals a transitional profibrotic macrophage. Nat. Immunol. 20, 163–172 (2019).

59. Kiselev, V. Y., Yiu, A. & Hemberg, M. scmap: projection of single-cell RNA-seq data across data sets. Nat. Methods 15, 359–362 (2018).

60. Zhang, A. W. et al. Probabilistic cell-type assignment of single-cell RNA-seq for tumor microenvironment profiling. Nat. Methods 16, 1007–1015 (2019).

61. Cao, Y., Wang, X. & Peng, G. SCSA: A Cell Type Annotation Tool for Single-Cell RNA-seq Data. Front. Genet. 11, (2020).

62. Tan, Y. & Cahan, P. SingleCellNet: A Computational Tool to Classify Single Cell RNA-Seq Data Across Platforms and Across Species. Cell Syst. 9, 207–213.e2 (2019).

63. Chen, J. et al. Transformer for one stop interpretable cell type annotation. Nat. Commun. 14, 223 (2023).

64. Franzén, O., Gan, L.-M. & Björkegren, J. L. M. PanglaoDB: a web server for exploration of mouse and human single-cell RNA sequencing data. Database 2019, baz046 (2019).

65. Smedley, D. et al. BioMart – biological queries made easy. BMC Genomics 10, 22 (2009).

66. Acland, A. et al. Database resources of the National Center for Biotechnology Information. Nucleic Acids Res. 42, D7–D17 (2014).

67. Alayoubi, Y., Bentsen, M. & Looso, M. Scanpro is a tool for robust proportion analysis of single-cell resolution data. Sci. Rep. 14, 15581 (2024).

68. Saxena, S. et al. Role of annexin gene and its regulation during zebrafish caudal fin regeneration. Wound Repair Regen. 24, 551–559 (2016).

69. Liu, W. & Hajjar, K. A. The annexin A2 system and angiogenesis. Biol. Chem. 397, 1005–1016 (2016).

70. Shi, S. et al. FGF19 promotes nasopharyngeal carcinoma progression by inducing angiogenesis via inhibiting TRIM21-mediated ANXA2 ubiquitination. Cell. Oncol. 47, 283–301 (2024).

71. Partevian, S. A. et al. The Effect of Morpholino Oligonucleotides to Gene Anxa2a on the Embryonic Development of Danio rerio. Mol. Genet. Microbiol. Virol. 38, 143–149 (2023).

72. Zhang, M. et al. Selenomethionine promotes ANXA2 phosphorylation for proliferation and protein synthesis of myoblasts and skeletal muscle growth. J. Nutr. Biochem. 115, 109277 (2023).

73. Li, J. et al. Grouper annexin A2 affects RGNNV by regulating the host immune response. Fish Shellfish Immunol. 137, 108771 (2023).

74. Bharadwaj, A., Kempster, E. & Waisman, D. M. The Annexin A2/S100A10 Complex: The Mutualistic Symbiosis of Two Distinct Proteins. Biomolecules 11, 1849 (2021).

75. Cmoch, A., Strzelecka-Kiliszek, A., Palczewska, M., Groves, P. & Pikula, S. Matrix vesicles isolated from mineralization-competent Saos-2 cells are selectively enriched with annexins and S100 proteins. Biochem. Biophys. Res. Commun. 412, 683–687 (2011).

76. 10k Human PBMCs, ATAC v2, Chromium Controller. 10x Genomics https://www.10xgenomics.com/datasets/10k-human-pbmcs-atac-v2-chromium-controller-2-standard.

77. Detleffsen, J., Bruns, B., Bentsen, M., Kuenne, C. & Looso, M. PEAKQC: periodicity evaluation in single-cell ATAC-seq data for quality assessment. Brief. Bioinform. 26, bbaf465 (2025).

78. Kondili, M. et al. UROPA: a tool for Universal RObust Peak Annotation. Sci. Rep. 7, 2593 (2017).

79. Kwok, A. W. C., Shim, H. & McCarthy, D. J. Going beyond cell clustering and feature aggregation: Is there single cell level information in single-cell ATAC-seq data? 2024.12.04.626927 Preprint at 10.1101/2024.12.04.626927 (2024).

80. Martens, L. D., Fischer, D. S., Yépez, V. A., Theis, F. J. & Gagneur, J. Modeling fragment counts improves single-cell ATAC-seq analysis. Nat. Methods 21, 28–31 (2024).

81. Cusanovich, D. A. et al. The cis-regulatory dynamics of embryonic development at single-cell resolution. Nature 555, 538–542 (2018).

82. Kleiveland, C. R. Peripheral Blood Mononuclear Cells. in The Impact of Food Bioactives on Health: in vitro and ex vivo models (eds Verhoeckx, K. et al.) (Springer, Cham (CH), 2015).

83. Peripheral Blood | Whole blood | Handbook | Miltenyi Biotec | Deutschland. https://www.miltenyibiotec.com/DE-en/support/macs-handbook/human-cells-and-organs/human-cell-sources/blood-human.html.

84. Bentsen, M. et al. ATAC-seq footprinting unravels kinetics of transcription factor binding during zygotic genome activation. Nat. Commun. 11, 4267 (2020).

85. Zwick, D., Vo, M. T., Shim, Y. J., Reijonen, H. & Do, J. BACH2: The Future of Induced T-Regulatory Cell Therapies. Cells 13, 891 (2024).

86. Titcombe, P. J., Silva Morales, M., Zhang, N. & Mueller, D. L. BATF represses BIM to sustain tolerant T cells in the periphery. J. Exp. Med. 220, e20230183 (2023).

87. Patterson, A. M. et al. Lung-resident memory CD4+ T cells are dependent on Batf3. J. Immunol. 214, 1133–1140 (2025).

88. Wang, B. & Bian, Q. SATB1 prevents immune cell infiltration by regulating chromatin organization and gene expression of a chemokine gene cluster in T cells. *Commun*. Biol. 7, 1304 (2024).

89. Gupta, P. K. et al. Reduced Satb1 expression predisposes CD4+ T conventional cells to Treg suppression and promotes transplant survival. Proc. Natl. Acad. Sci. U. S. A. 119, e2205062119 (2022).

90. Kim, H. S., Sohn, H., Jang, S. W. & Lee, G. R. The transcription factor NFIL3 controls regulatory T-cell function and stability. Exp. Mol. Med. 51, 1–15 (2019).

91. Yang, X. et al. ATF4 Regulates CD4+ T Cell Immune Responses through Metabolic Reprogramming. Cell Rep. 23, 1754–1766 (2018).

92. Wei, X. et al. The Multifaceted Roles of BACH1 in Disease: Implications for Biological Functions and Therapeutic Applications. Adv. Sci. 12, 2412850 (2025).

93. Song, J. et al. Charged substrate treatment enhances T cell mediated cancer immunotherapy. Nat. Commun. 16, 1585 (2025).

94. Wagle, M. V. et al. Antigen-driven EGR2 expression is required for exhausted CD8+ T cell stability and maintenance. Nat. Commun. 12, 2782 (2021).

95. Fagerberg, E. et al. KLF2 maintains lineage fidelity and suppresses CD8 T cell exhaustion during acute LCMV infection. Science 387, eadn2337 (2025).

96. Lacorazza, H. D. The reprogramming factor KLF4 in normal and malignant blood cells. Front. Immunol. 16, (2025).

97. Moskowitz, D. M. et al. Epigenomics of human CD8 T cell differentiation and aging. Sci. Immunol. 2, eaag0192 (2017).

98. Zhang, H., et al. Sustained AhR activity programs memory fate of early effector CD8+ T cells. Proc. Natl. Acad. Sci. 121, e2317658121 (2024).

99. Tombor, L. S. et al. Single cell sequencing reveals endothelial plasticity with transient mesenchymal activation after myocardial infarction. Nat. Commun. 12, 681 (2021).

100. Khassafi, F. et al. Transcriptional profiling unveils molecular subgroups of adaptive and maladaptive right ventricular remodeling in pulmonary hypertension. *Nat*. Cardiovasc. Res. 2, 917–936 (2023).

101. Dizdarevic, S. et al. Mapping the Immune Landscape of Lung Cancer: Identification of Unique CD45+ Cell Subsets. Am. J. Respir. Crit. Care Med. 211, A3098–A3098 (2025).

102. Panza, P. et al. The lung microvasculature promotes alveolar type 2 cell differentiation via secreted SPARCL1. Stem Cell Rep. 20, 102451 (2025).

103. Xu, Y. et al. PDGFRA is a conserved HAND2 effector during early cardiac development. *Nat*. Cardiovasc. Res. 3, 1531–1548 (2024).

104. Wang, Z.-Y. et al. flt1 inactivation promotes zebrafish cardiac regeneration by enhancing endothelial activity and limiting the fibrotic response. Development 151, dev203028 (2024).

105. Goumenaki, P. et al. The innate immune regulator MyD88 dampens fibrosis during zebrafish heart regeneration. *Nat*. Cardiovasc. Res. 3, 1158–1176 (2024).

106. Malainou, C. et al. TNF Superfamily Member 14 Drives Post-Influenza Depletion of Alveolar Macrophages Enabling Secondary Pneumococcal Pneumonia. 2024.07.28.605445 Preprint at 10.1101/2024.07.28.605445 (2024).

107. Jang, J. et al. Endocardial HDAC3 is required for myocardial trabeculation. Nat. Commun. 15, 4166 (2024).

108. da Silva, A. R. et al. egr3 is a mechanosensitive transcription factor gene required for cardiac valve morphogenesis. Sci. Adv. 10, eadl0633 (2024).

109. Cardeira-da-Silva, J. et al. Antigen presentation plays positive roles in the regenerative response to cardiac injury in zebrafish. Nat. Commun. 15, 3637 (2024).

110. Juan, T. et al. Control of cardiac contractions using Cre-lox and degron strategies in zebrafish. Proc. Natl. Acad. Sci. 121, e2309842121 (2024).

111. Roquid, K. A. et al. Lung endothelial PEAR1 induces tumor cell dormancy. Mol. Cancer 24, 278 (2025).

112. Chu, X. et al. GLI1+ Cells Contribute to Vascular Remodeling in Pulmonary Hypertension. Circ. Res. 134, e133–e149 (2024).

113. Dimitrov, D. et al. LIANA+ provides an all-in-one framework for cell–cell communication inference. Nat. Cell Biol. 26, 1613–1622 (2024).

114. Raredon, M. S. B. et al. Computation and visualization of cell–cell signaling topologies in single-cell systems data using Connectome. Sci. Rep. 12, 4187 (2022).

