## Supplementary Figures for "SC-Framework: A robust and FAIR semi-interactive environment for single-cell resolution datasets"

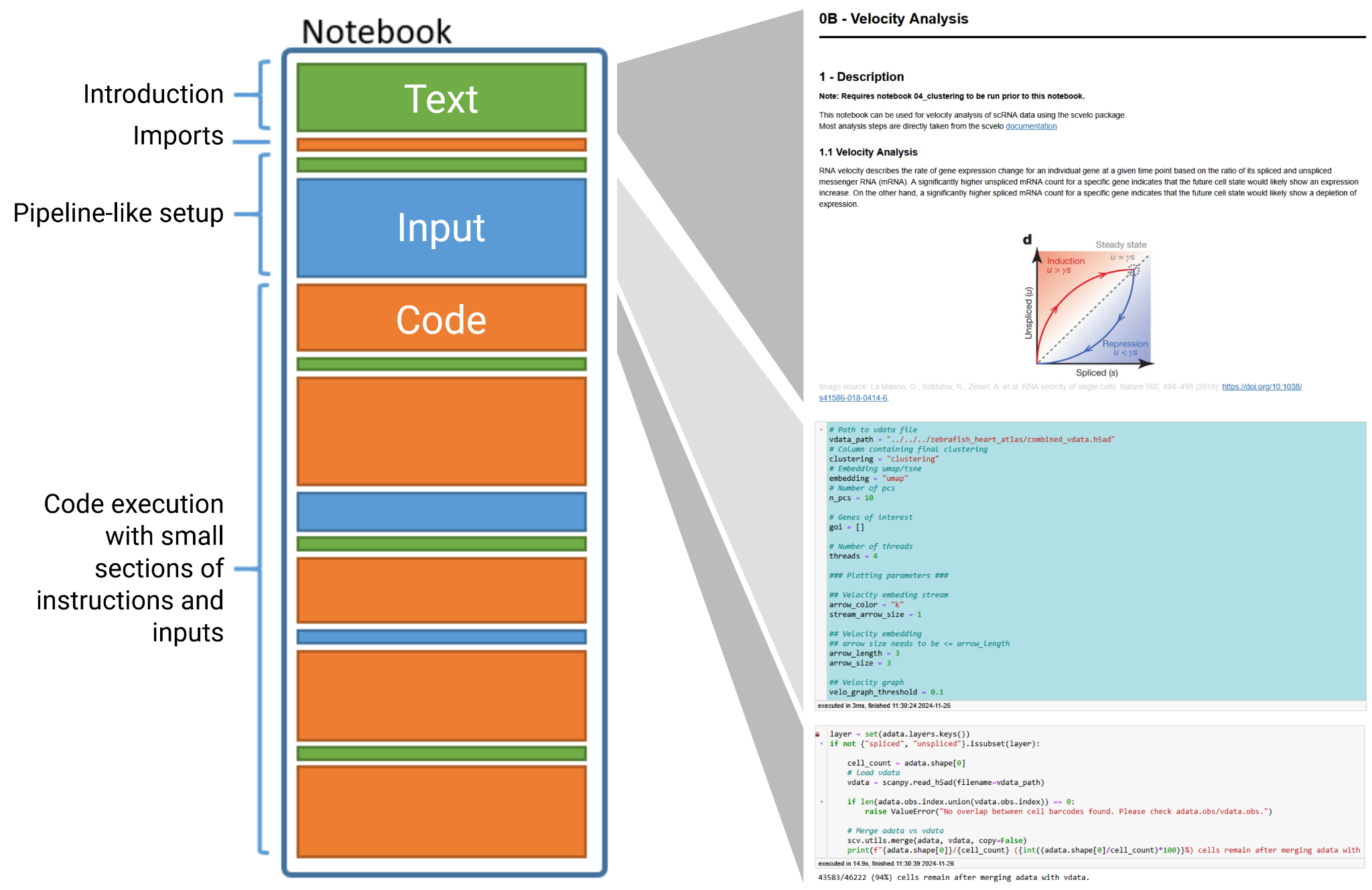

**Supp. Figure 1: Notebook structure**

The general structure that each notebook follows (left) with excerpts from the velocity notebook (right). All notebooks start with an introduction text (green) providing an overview of the specific notebook's use case. This is followed by a small code block to load the functions required for the following steps (orange). Cells with a blue background indicate user intervention; the first blue cell provides global parameters for the notebook. Text and input cells (green and blue) later in the notebook describe specific analysis steps and guide the user to set optimal parameters.

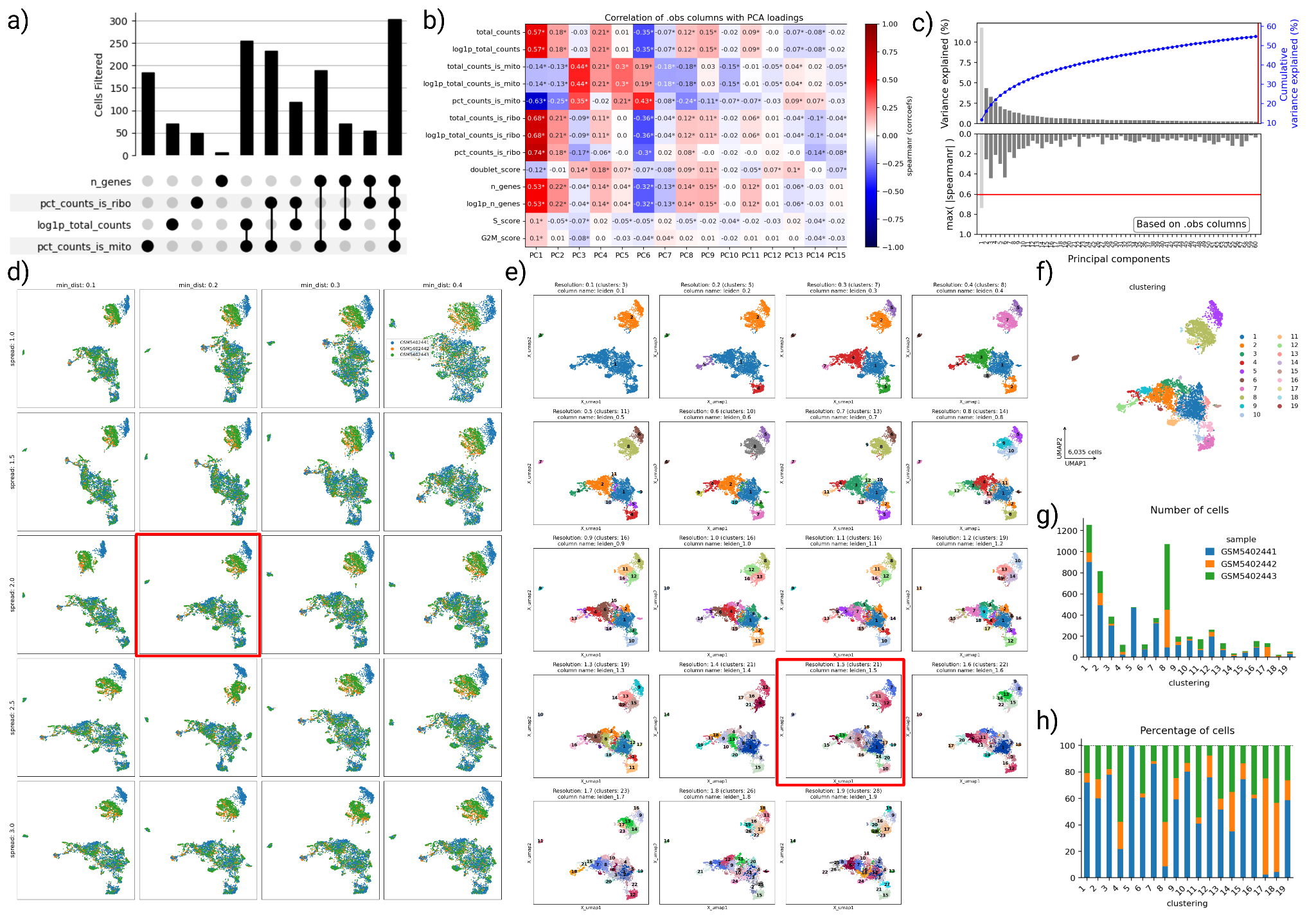

**Supp. Figure 2: zebrafish 150 dpf core analysis plots**

a) an UpSet plot showing the impact each metric (alone and combined) has during cell filtering. b) a heatmap of filter metrics (rows) and PCs (columns), colored for the correlation of the metric to the respective PC. Cropped to the first 15 PCs. c) barplots with suggested PC filters. The top barplot shows the explained variance of each PC (bar), with a threshold set to keep all PCs. The bottom barplot shows the maximum absolute correlation (see b) for each PC. Here, the first bar is light grey because it exceeds the correlation threshold and is therefore suggested for removal. d) grid of UMAP embeddings with different parameter combinations to find the optimal parameters, with the highlighted UMAP selected for further analysis. e) same as in d) but for optimizing the clustering. f) final UMAP and clustering after selecting and further adjusting UMAP and clustering of d) and e). g-h) barplot of sample distribution in each cluster, with raw cell counts (g) and percentage (h).

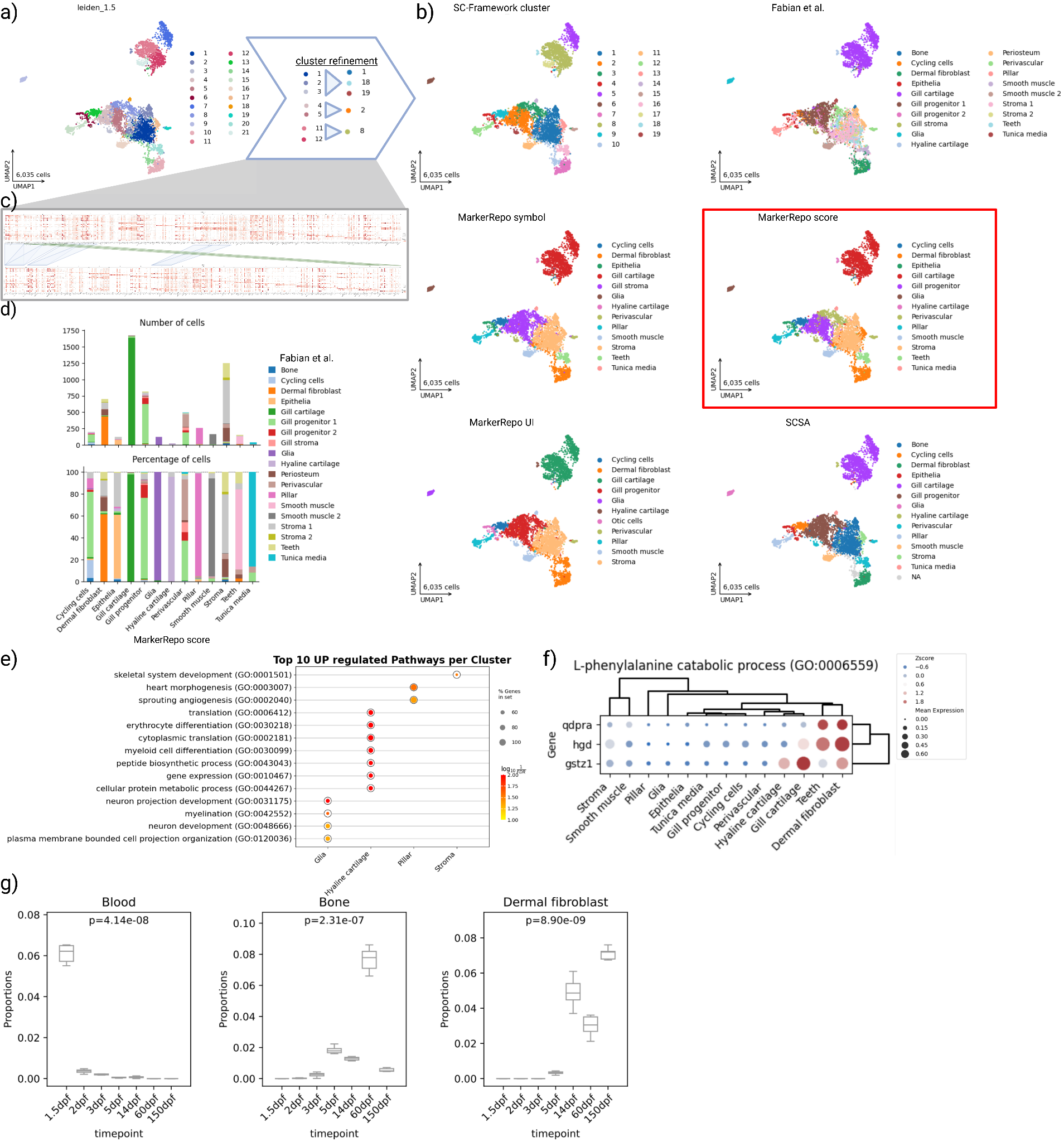

**Supp. Figure 3: downstream plots of zebrafish 150 dpf and combined timepoints**

a) UMAP embedding colored for the initially chosen clustering and cluster adjustments based on marker genes to the right. b) UMAP with final clustering after adjustments (top-left), reference annotation of Fabian et al. (top-right), and MarkerRepo predictions based on different algorithms. The marked annotation is chosen for the following analysis b). Dot plots showing the marker genes before (top) and after (bottom) cluster adjustments c). Barplot of MarkerRepo annotated cell types d) compared to the Fabian et al. annotation. e) GO Term enrichment analysis showing the top significant terms per cell type. f) Expression of phenylalanine catabolic process genes in cell types. g) Changes in cell proportion over time for selected cell types. Each plot shows the proportion of cells (y-axis) assigned to the respective cell type per time point (x-axis). A p-value provides the significance of the proportional changes in cell type composition. Created in BioRender. Schultheis, H. (2025) https://BioRender.com/lqmg4vy

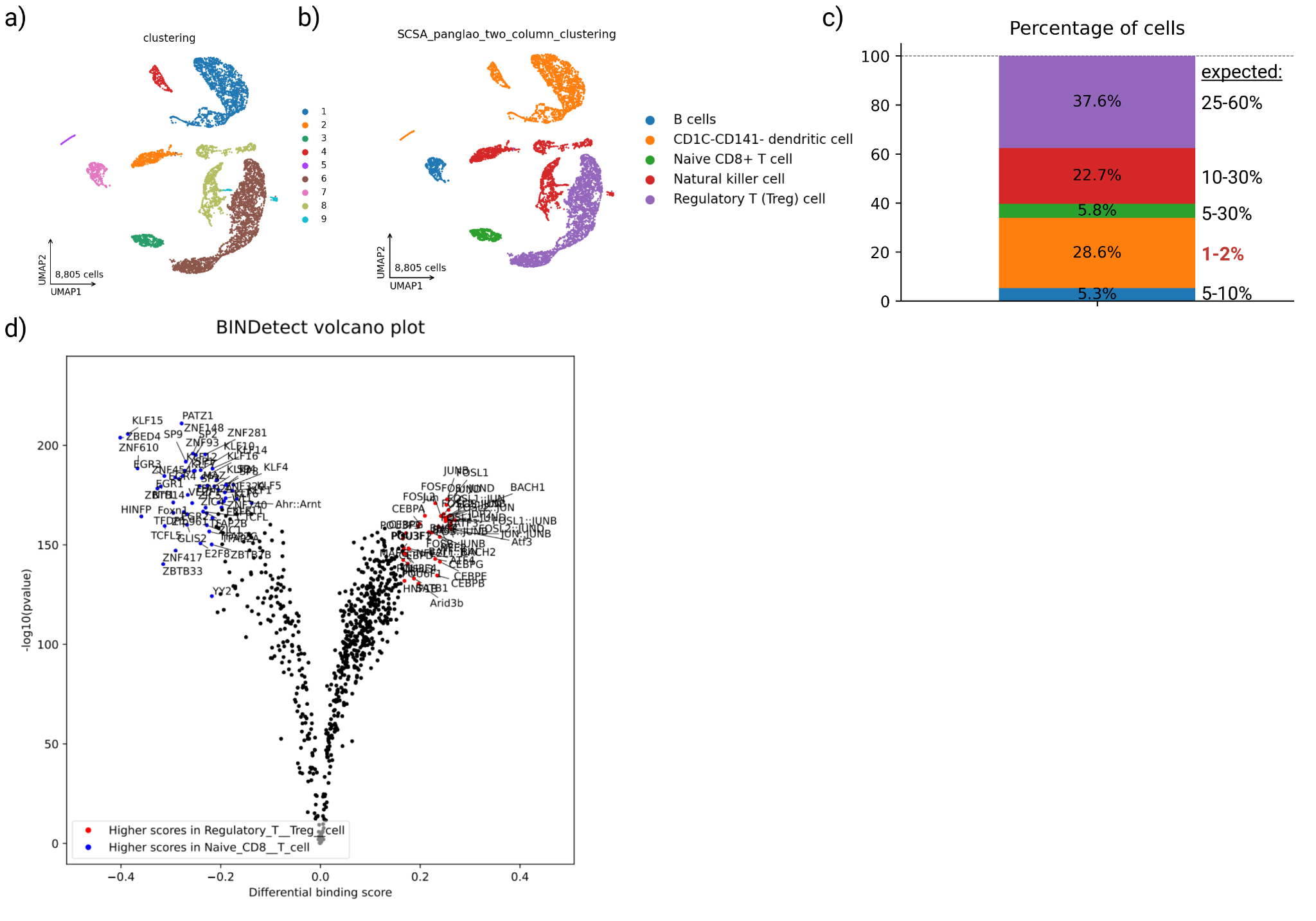

**Supp. Figure 4: Key aspects of the PBMC snATAC analysis.**

UMAP embedding and initial division into nine clusters (a). The UMAP embedding colored by five PBMC-related cell types, predicted by our MarkerRepo tool (b). The percentage of cells for each predicted cell type and expected percentages (MACS handbook) (c). A TOBIAS BINDetect volcano plot, showing differential transcription factor binding. Top transcription factors enriched for Naive CD8^+^ T cells are shown in blue, and top enriched transcription factors in Regulatory T cells are shown in red.
